# DNA transposon expansion is associated with genome size increase in mudminnows

**DOI:** 10.1101/2021.06.07.447207

**Authors:** Robert Lehmann, Aleš Kovařík, Konrad Ocalewicz, Lech Kirtiklis, Andrea Zuccolo, Jesper N. Tegner, Josef Wanzenböck, Louis Bernatchez, Dunja K. Lamatsch, Radka Symonová

**Affiliations:** Division of Biological and Environmental Sciences & Engineering, Computer, Electrical and Mathematical Sciences and Engineering Division, King Abdullah University of Science and Technology, Thuwal, 23955-6900, Kingdom of Saudi Arabia; Laboratory of Molecular Epigenetics, Institute of Biophysics, Czech Academy of Science, Královopolská 135, 61265, Brno, Czech Republic; Department of Marine Biology and Ecology, Institute of Oceanography, Faculty of Oceanography and Geography, University of Gdansk, Gdansk, Poland; Department of Zoology, Faculty of Biology and Biotechnology, University of Warmia and Mazury, M. Oczapowskiego Str. 5, 10-718, Olsztyn, Poland; Center for Desert Agriculture, Biological and Environmental Sciences & Engineering Division (BESE), King Abdullah University of Science and Technology, Thuwal 23955-6900, Saudi Arabia; Institute of Life Sciences, Scuola Superiore Sant’Anna, Pisa 56127, Italy; Research Department for Limnology Mondsee, University of Innsbruck, A – 5310 Mondsee, Austria; IBIS (Institut de Biologie Intégrative et des Systèmes), Université Laval, Québec, QC, Canada; Department of Bioinformatics, Wissenschaftzentrum Weihenstephan, Technische Universität München, Freising, Germany; Faculty of Biology, University of Hradec Kralove, Czech Republic

**Keywords:** Genome expansion, *Umbra*, Robertsonian fusion, centric fission, repetitive sequences

## Abstract

Genome sizes of eukaryotic organisms vary substantially, with whole genome duplications (WGD) and transposable element expansion acting as main drivers for rapid genome size increase. The two North American mudminnows, *Umbra limi* and *U. pygmaea*, feature genomes about twice the size of their sister lineage Esocidae (e.g., pikes and pickerels). However, it is unknown whether all *Umbra* species share this genome expansion and which causal mechanisms drive this expansion. Using flow cytometry, we find that the genome of the European mudminnow is expanded similarly to both North American species, ranging between 4.5-5.4 pg per diploid nucleus. Observed blocks of interstitially located telomeric repeats in *Umbra limi* suggest frequent Robertsonian rearrangements in its history. Comparative analyses of transcriptome and genome assemblies show that the genome expansion in *Umbra* is driven by extensive DNA transposon expansion without WGD. Furthermore, we find a substantial ongoing expansion of repeat sequences in the Alaska blackfish *Dallia pectoralis*, the closest relative to the family Umbridae, which might mark the beginning of a similar genome expansion. Our study suggests that the genome expansion in mudminnows, driven mainly by transposon expansion, but not WGD, occurred before the separation into the American and European lineage.

**Significance Statement:** North American mudminnows feature genomes about twice the size of their sister lineage Esocidae (e.g., pikes and pickerels). However, neither the mechanism underlaying this genome expansion, nor whether this feature is shared amongst all mudminnows is currently known. Using cytogenetic analyses, we find that the genome of the European mudminnow also expanded and that extensive chromosome fusion events have occurred in some Umbra species. Furthermore, comparative genomics based on de-novo assembled transcriptomes and genome assemblies, which have recently become available, indicates that DNA transposon activity is responsible for this expansion.

## Introduction

The driving forces and effects of genome size variations across different taxa are a recurring theme in the field of evolutionary biology (Lynch 2007). While the number of chromosomes is typically highly conserved among teleost fishes (Mank & Avise 2006), genome sizes vary substantially and include the smallest and the largest vertebrate genome (Hardie & Hebert 2004). Specifically, fish genome size ranges from C = 0.35 pg in bandtail pufferfish (*Sphoeroides spengleri)* to C = 133 pg in marbled lungfish (*Protopterus aethiopicus)* (Gregory 2020). The main candidate processes introducing such drastic variation are gene-(Ohno 2013; Lu et al. 2012) or genome duplication(Fontdevila 2011; Van de Peer et al. 2017), transposable element (TE) proliferation (Pritham 2009; Tenaillon et al. 2010), and replication slippage at tandem repeat loci(Ellegren 2004). Furthermore, it is unclear whether this variation is shaped by adaptive processes (Gregory & Hebert 1999; Liedtke et al. 2018; Van de Peer et al. 2017) or stochastic sequence gain and loss(Lynch 2007). The presence of TEs in virtually all eukaryotic genomes suggests a role of stochastic sequence gain due to TE proliferation (Elliott & Gregory 2015). There are several recent reports of significant genomic expansion, such as in salamanders due to long terminal repeat element activity (Sun et al. 2012) and in *Hydra* due to long interspersed nuclear element activity (Wong et al. 2019). Estimates of the pervasiveness and extent of expansion events caused by TE proliferation and other processes will provide insight into the forces that shape genome size evolution.

Mudminnows (Umbridae) are very resilient members of the order Esociformes, capable of withstanding extreme cold and can utilize atmospheric oxygen. Under adverse conditions, they are reputed to become dormant in mud (Gilbert & Williams 2002). The order Esociformes includes two families and four genera in at least 12 species with one fossil-only family (Nelson et al. 2016). Furthermore, Esociformes (Haplomi, Esocae; pikes and mudminnows) and its sister order Salmoniformes belong to the clade Protacanthopterygii, a lineage of basal teleosts (Nelson et al. 2016). Genome sizes of both North American *Umbra* species were determined already in the 60s to be C = 2.52 - 2.70 pg for *Umbra limi* and C = 2.4 pg for *U. pygmaea* (Hinegardner 1968). The genome size of the remaining representatives of the order Esociformes, including *Esox*, *Novumbra*, and *Dallia*, ranges from 0.85 (in *Esox*) to 1.4 (also in *Esox*; summarized by (Gregory 2020); details in Table 1). While two of the three extant *Umbra* species feature a genome size twice the size of the remaining esociform species, it is currently not known whether this is a universal feature of the Umbridae and can be expected in *U. krameri*.

**Table 1.** Cytogenomic data for extant species of the order Esociformes from http://www.genomesize.com, http://www.animalrdnadatabase.com, literature, and results obtained in this study.

| Family | Species | C-val (pg) | 2n/FN | Method | 45S rDNA | 5S rDNA | Reference |
| --- | --- | --- | --- | --- | --- | --- | --- |
| Esocidae | *E. americanus americanus* | 1.18 | 50/72 | FD |  |  | (Beamish et al. 1971; Rab & Crossman 1994) |
| Esocidae | *E. americanus vermiculatus* | 1.06/1.12 | 50 | FIA, FD |  |  | (Beamish et al. 1971; Hardie & Hebert 2004) |
| Esocidae | *E. lucius* | 0.97 |  | BCA |  |  | (Vendrely & Vendrely 1953, 1952a) |
| Esocidae | *E. lucius* | 0.98 |  | BCA |  |  | (Vendrely & Vendrely 1952b) |
| Esocidae | *E. lucius* | 1.13 |  | FIA |  |  | (Hardie & Hebert 2003, 2004) |
| Esocidae | *E. lucius* | 1.15 | 50/<br>50-58 | FCM |  |  | (Vinogradov 1998;<br>Khoshkholgh et al. 2015) |
| Esocidae | *E. lucius* | 1.36 |  | FD |  |  | (Beamish et al. 1971) |
| Esocidae | *E. lucius* | 1.11 | 50/50 | FCM | 2 | 30-38 | present study;<br>(Symonová et al. 2017) |
| Esocidae | *E. cisalpinus* |  | 50/50 | N/A | 2 | 30-34 | (Symonová et al. 2017) |
| Esocidae | *E. masquinongy* | 1.28 | 50/<br>76-78 | FD | 2 |  | (Beamish et al. 1971; Rab &<br>Crossman 1994) |
| Esocidae | *E. niger* | 0.92 |  | FIA |  | 4 | (Hardie & Hebert 2003, 2004);<br>present study |
| Esocidae | *E. niger* | 1.18 | 50/74 | FD | 2 |  | (Beamish et al. 1971) |
| Esocidae | *E. reichertii* | 1.28 | 50 | FD |  |  | (Beamish et al. 1971) |
| Esocidae | *Dallia pectoralis* | 1.26 | 74-78/ | FD | 6 |  | (Beamish et al. 1971; Rab &<br>Crossman 1994) |
| Esocidae | *Novumbra hubbsi* | 1.04 | 48/62 | FD | 12 |  | (Beamish et al. 1971; Crossman<br>& Ráb 2001) |
| Umbridae | *U. limi* | 2.52 |  | FD |  | 3 | (Beamish et al. 1971); present<br>study |
| Umbridae | *U. limi* | 2.56 |  | FIA |  |  | (Hardie & Hebert 2004) |
| Umbridae | *U. limi* | 2.70 | 22/44 | BFA | 2 |  | (Hinegardner 1968; Hinegardner<br>& Rosen 1972) |
| Umbridae | *U. krameri* | 2.27 | 44/44 | FCM |  |  | present study; reviewed in (Arai<br>2011) |
| Umbridae | *U. pygmaea* | 2.41 | 22/44 | FD | 2-4 |  | (Beamish et al. 1971) |

Abbreviations used: Method: BCA – biochemical analysis, BFA – Bulk fluorometric assay, FD – Feulgen densitometry, FIA - Feulgen image analysis densitometry, FCM – Flow cytometry, N/A – not available.

The diploid chromosome number in Esociformes ranges from 2n = 22 (*U. pygmaea* and *U. limi*) to 2n = 71 - 79 (*Dallia* sp.). Almost all known pike (*Esox*) species possess 50 (fundamental number FN = 50) usually acrocentric chromosomes, including species from Canada, USA, Sweden, and Russia (Arai 2011) and two European *Esox* species (Symonová et al. 2017). Such karyotype is considered the pre-duplication ancestor of the order Salmoniformes (salmon, grayling, whitefish) (Rondeau et al. 2014; Ráb 2004). There are, however, reports of FN = 86 with 2n = 50 in *E. lucius* from the south Caspian Sea basin and further reports of FN = 72 - 88 in other *E. lucius* populations (Khoshkholgh et al. 2015). This morphological variability of chromosomes in the *Esox* genus with its broad circumpolar distribution (Nelson et al. 2016) has not yet been analyzed.

Pikes are highly interesting from the cytogenomic viewpoint because of their extremely amplified, massively methylated, and potentially functional 5S rDNA recorded on more than 30 centromeric sites of their 50 chromosomes, whereas 45S rDNA is located in only two sites (Symonová et al. 2017). A similar amplification and increased dynamics of the 45S rDNA fraction has been repeatedly recorded in Salmoniformes ^12,13^, which furthermore underwent an already well-characterized whole-genome duplication (WGD) (Macqueen & Johnston 2014; Lien et al. 2016). Indeed, WGD events have been described for multiple basal, particularly freshwater teleosts such as Cypriniformes (Li et al. 2015) and Siluriformes (Marburger et al. 2018). This observation suggests two distinct explanations for the significant genome size increase of Umbridae compared to its sister lineage Esocidae, an Umbridae-specific WGD on the one hand, and an extreme amplification of ribosomal DNA on the other.

Here, we present cytogenomic evidence that all extant members of the *Umbra* lineage feature an expanded genome size compared to its closest relative, the Esocidae family. Furthermore, we find that Umbridae feature a standard locus and copy number of 5S rDNA typical for teleosts, excluding a 5S rDNA expansion as a reason for the *Umbra*-specific genome. While phylogenetic profiles of orthologous genes across species do not show patterns of WGD, comparative analyses of repetitive sequence content across six Esocidae species show a clear signal of DNA transposon expansion in *Umbra pygmaea* driving the genome size increase.

## Results

### Flow cytometry genome size determination in the European pike and the European mudminnow

Flow cytometry measurements using DAPI staining of *U. krameri* with chicken RBC as internal standard exhibited a genome size of 2C = 4.54 ± 0.05 pg (±STDEV) per nucleus (CV% 3.36 ± 0.55; range 4.43-4.60 pg per nucleus, N=10). *E. lucius* gave genome values of 2C = 2.22 ± 0.03 pg per nucleus (CV% 2.44 ± 1.37, range 2.18 - 2.27 pg per nucleus, N=2). Genome size and chromosome traits (2n and FN) for the order Esociformes are summarized in Fig. 1 and show the elevated genome size in all three *Umbra* species. The higher interquartile range in *Esox lucius* reflects merely the higher sampling effort and multiple values available. A similar situation exists in *Umbra limi*. We provide an overview of cytogenomic and cytogenetic traits of the order Esociformes (Table 1) based on the checklist of fish karyotypes (Arai 2011).

**Fig.1.**
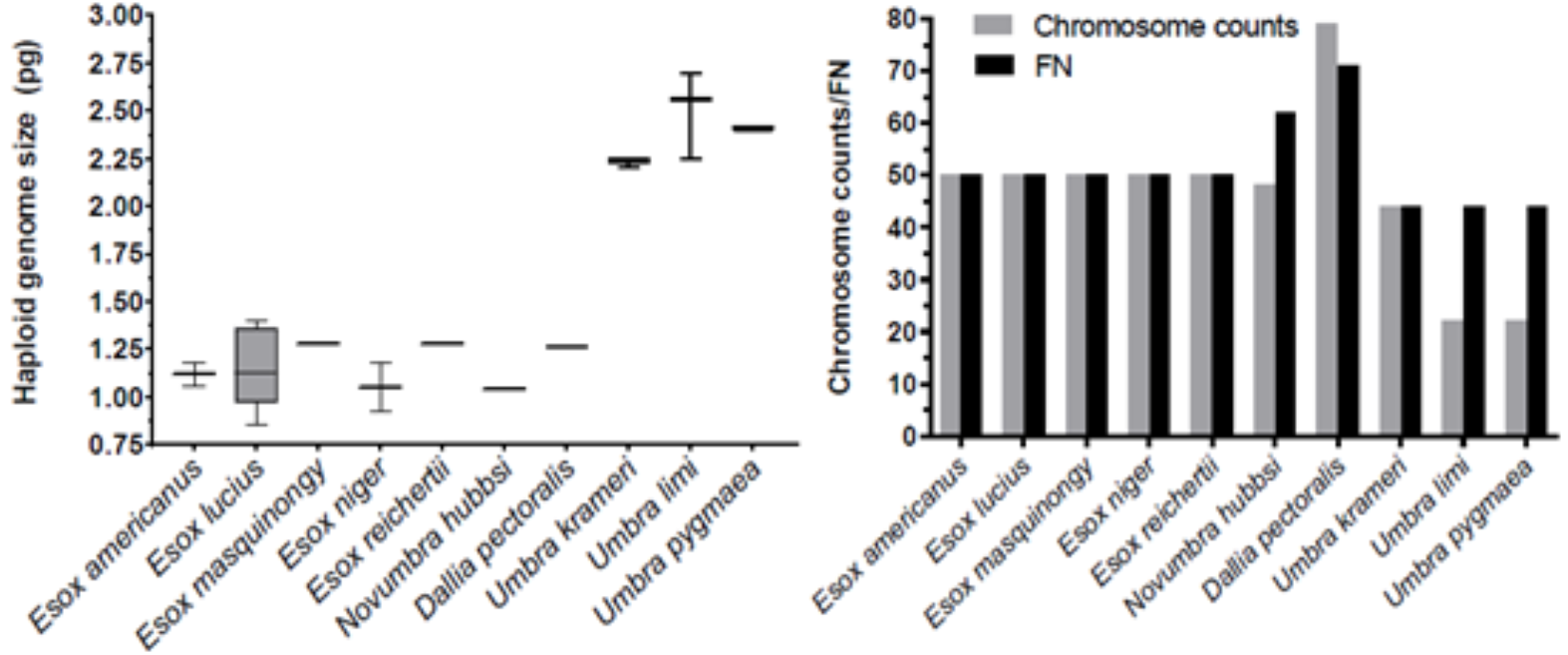
Genome size (C-value) (left) and chromosome (2n) and chromosome arms (FN) counts (right) in esociform fish.

### Comparative analysis of the *U. pygmaea* transcriptome does not indicate whole genome duplication

To test whether the *Umbridae* family underwent a whole-genome duplication event (WGD), we assessed gene duplication levels in a set of 24 *de novo* transcriptome assemblies, including the eastern mudminnow *U. pygmaea* (Pasquier et al. 2016). We focused on a set of 3640 benchmark universal single-copy orthologous (BUSCO) genes, which occur only once in the majority (>90%) of a set of 26 selected Actinopterygii species (Simão et al. 2015). Since only a single salmonid species, *Salmo salar,* was used to construct the BUSCO set, genes that were duplicated in the salmonid-specific WGD(Macqueen & Johnston 2014) were not excluded. Similarly, as this gene set does not consider *Umbridae* family species, any potential ohnologs originating from an *Umbridae*-specific WGD are not excluded. Accordingly, five of the six salmonid *de novo* transcriptome assemblies show the highest duplication level amongst the 24 considered species (Fig. S2, Table S1). In contrast, *U. pygmaea* exhibits the second-lowest duplication level, second only to *Pangasius hypophthalmus*. The elevated duplication level in European eel *Anguilla anguilla* and arowana *Osteoglossum bichirrosum* were noted recently, suggesting an additional WGD as a possible cause(Rozenfeld et al. 2019).

Transcriptome d*e novo* assembly procedures include steps that can influence the observed number of duplicated genes, such as the clustering of the assembled contigs(Pasquier et al. 2016; Grabherr et al. 2011). Therefore, we collected all available reference genome assemblies matching the species of any of the transcriptome assemblies. The direct comparison of BUSCO estimates for *de novo* assemblies and reference genome-derived transcriptomes across 10 species revealed a high correspondence of the duplication level (R^2^ = 0.8) (Fig. S1, Table S2). In contrast, parameters reflecting technical differences of transcriptome generation show substantially lower correspondence, such as the number of fragmented (R^2^ = 0.02) or missing transcripts (R^2^ = 0.28). Clustering of BUSCO gene phylogenetic profiles across all considered species provides a more detailed picture (Fig. 2). While a small group of 240 transcripts fails to assemble reliably (cluster I), likely reflecting specific sequence properties, the second group of 819 genes is detected as single-copy in the majority of species (cluster II). The largest cluster III, comprised of 2096 genes, exhibits species-specific duplication patterns. Cluster IV contains 301 genes duplicated mainly in salmonids and thus likely represent remnants of the salmonid-specific WGD (cluster IV). The remaining 184 genes appear almost consistently duplicated or triplicated across all considered species, again likely reflecting sequence properties (cluster V). Both members of the Esocidae, the eastern mudminnow *U. pygmaea,* and the Northern pike *Esox lucius* are placed next to each other in the hierarchical species clustering. Separating cluster III into subclusters reveals that both the eastern mudminnow and the Northern pike feature only small non-overlapping groups of species-specific duplicated genes containing 47 and 76 genes, respectively (Fig. S3, Table S3).

**Fig.2.**
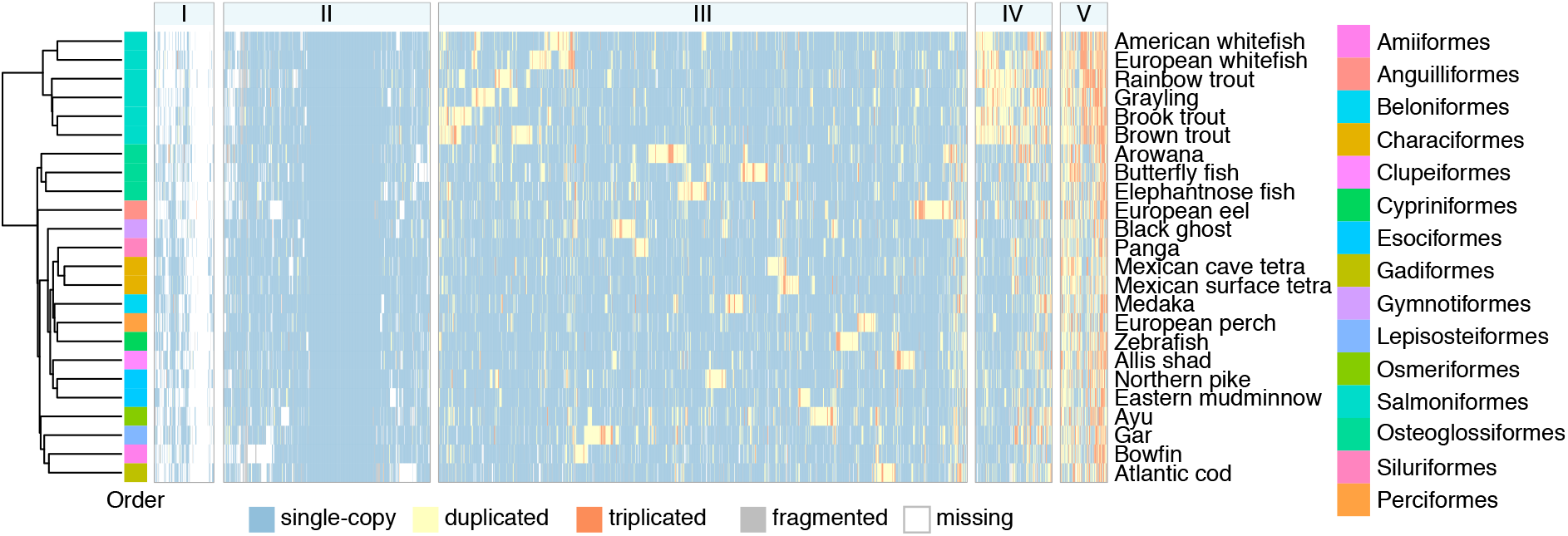
Phylogenetic profiling of universal single-copy orthologous genes in 24 *de novo* transcriptome assemblies. Genes are grouped into five clusters (top, I-V) by profile similarity. The hierarchical clustering tree for all species is shown on the left together with the corresponding color-coded order.

### Comparative genome analyses of Esociformes species reveal extensive transposable element expansions

We performed a direct comparative analysis of the genome composition of *U. pygmaea* using recently published short-read-based genome assemblies of five Esociformes species(Pan et al. 2021) and, in addition, the high-quality chromosome scale genome assembly of *E. lucius* (Table S4). Due to the employed short-read sequencing and relatively low coverages, the assemblies for *U. pygmaea*, *E. masquinongy*, *E. niger*, *N. hubbsi*, and *D. pectoralis* are fragmented and less complete (11 - 24% BUSCO genes missing) compared to the chromosome-scale assembly of *E. lucius* (3% missing). Nonetheless, the genome assembly sizes correspond well to direct measurements of genome size (Fig. 3A). In agreement with the transcriptome analysis, the assessment of BUSCO genes in all six genome assemblies shows low levels of duplication. We then characterized the repetitive sequence content of each genome aiming to test the hypothesis that transposable element activity might have caused a genome expansion exclusively in *U. pygmaea*. We, therefore, annotated repetitive sequences in all six genomes using a combined *de novo* repeat library (see Methods). The observed repeat content in each genome scales with the total genome size (Fig. 3B), with the highest repeat content of 64% in the largest genome of *U. pygmaea* and 37% of repeat content in the smallest genome of *N. hubbsi* (Fig. 3C, stacked plot on the left). In general, DNA transposons make up the largest part of classified repeat sequences in all analyzed species, contributing about 19% genomic sequence in *U. pygmaea*, *E. lucius*, and *E. masquinongy* (Tab. S4). In contrast, *E. niger*, *N. hubbsi*, and *D. pectoralis* show lower DNA transposon abundances with 14.8%, 10.8%, and 8%, respectively. A similar pattern is observed for LINE elements, while SINE elements and LTRs are generally low in abundance. A principal component analysis of repeat abundances across all six species reveals that the repetitive sequence composition mirrors the phylogenetic relationships (Fig. S4), with high similarity between *E. lucius* and *E. masquinongy*, as well as between *N. hubbsi* and *D. pectoralis*. Both, *U. pygmaea* and *E. niger,* constitute outliers in repeat composition. Importantly, we observe a significant expansion of highly repetitive and species-specific sequences awaiting further characterization in *U. pygmaea* with 30.8% genomic abundance, compared to abundances in the range of 15.6-21.2% in all other species.

**Fig.3.**
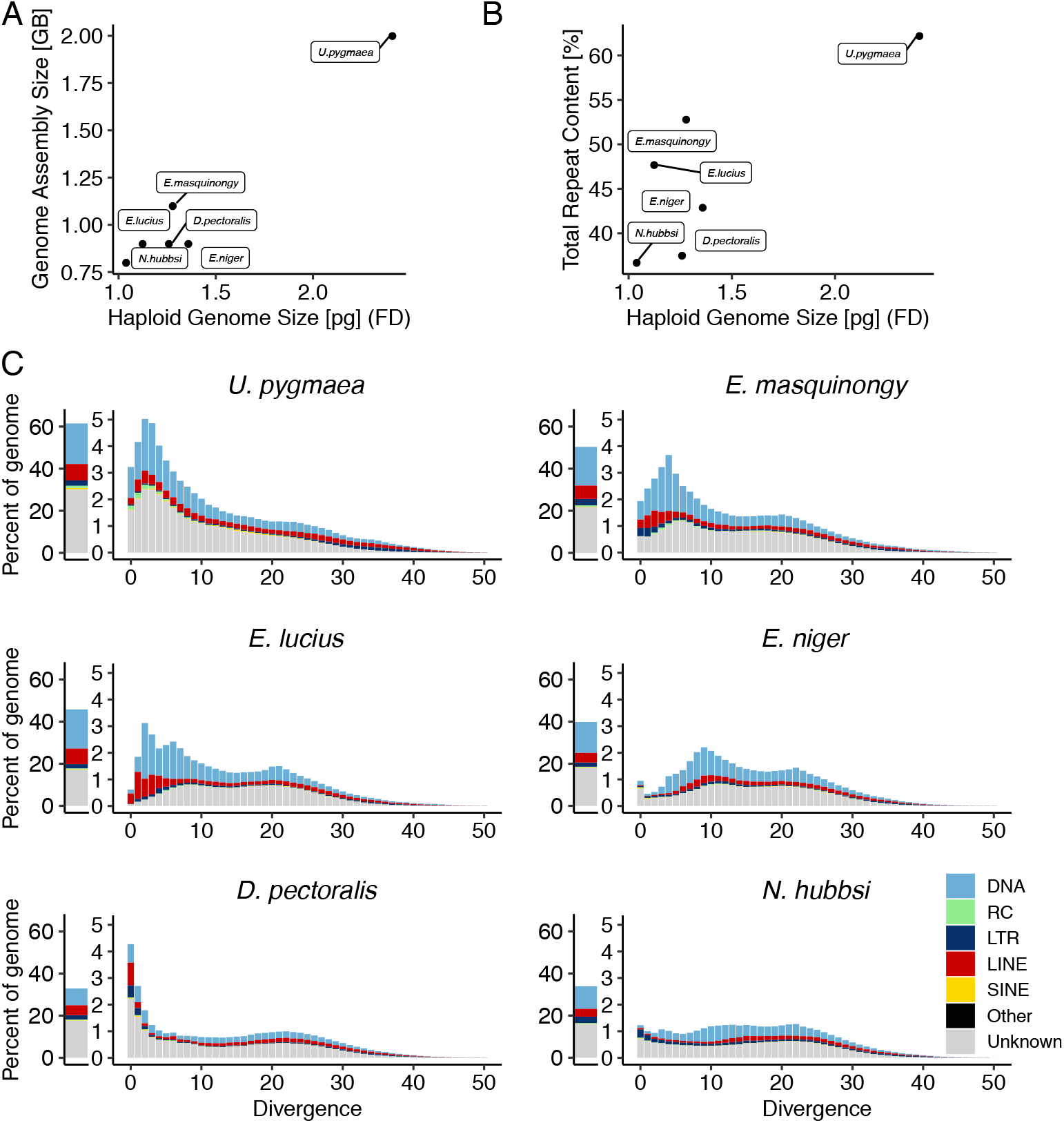
Genomic repeat sequence content and divergence landscapes in six Esociformes species. A) Genome size measurements by Feulgen densitometry compared to genome assembly size (Table S4). B) Genome assembly size compared to repetitive sequence content. C) Total repetitive sequence content (left) and repeat copy divergence landscapes (right) as fraction of the total genome assembly for each of the six species, distinguishing the main repetitive sequence classes.

Similar to the repeat composition, repeat expansion histories also show significant differences between the considered species (Fig. 3C). Here, the repeat expansion history is captured by the Kimura substitution level for each genomic copy of a repeat as a proxy for its age, since each copy accumulates mutations over time and diverges from its consensus sequence. The resulting divergence landscape suggests that the genome size increase of *U. pygmaea* results from a recent spike in DNA transposon activity combined with the accumulation of unclassified repetitive sequence, with 45% of the genomic repeat sequence showing a mean divergence below 8 (Fig. 3C, top left). The most abundant repeat sequence in *U. pygmaea* is the Tc1/mariner element IC-Mifl (upygm-1-8) which makes up 1% (20.5 Mb) of the genome assembly with an average divergence of 8 (Table S5). The second most abundant repeat Tc1-2_Xt (upygm-1-14), with 14 Mb and 0.7 % of genomic sequence, is also member of the Tc1/mariner family and shows a very recent expansion pattern with an average divergence of 1.34. *E. masquinongy* and *E. lucius* also show a notable recent expansion peak driven by DNA transposon activity. Similar to *U. pygmaea*, the expansion of *E. lucius* is also driven in part by IC-Mifl which contributes 1.3 % / 11.7 Mb to the total genome assembly, making it the third most abundant repeat only surpassed by Mariner-9 originally identified in *Salmo salar* and an RTE-1 non-long-terminal-repeat element identified in *E. lucius* (Table S5). The prominence of IC-Mifl notwithstanding, the high total abundance of DNA transposon sequence in the aforementioned species is caused by a multitude of active families. While the total repeat content of *D. pectoralis* is low, the divergence landscape reveals a currently ongoing repeat expansion with 27% of the total repeat content at divergences below 4, which translates into 9.1 % of the total genome sequence. In contrast to all other species, the expansion is caused by two LINE/RTE-BovB elements with high abundance (0.5 and 0.2%, respectively, of total genome) and low mean divergence (1.16 and 0.87) in combination with rRNA, tRNA, and unclassified sequences. This leaves *N. hubbsi* as the only species with a consistently low transposable element activity.

To gain a more detailed overview of the Tc1/mariner transposon superfamily activity, we constructed a phylogeny of all transposase sequences in our repeat library together with transposase sequences of a previously described Tc1/mariner reference set(Gao et al. 2017) in neoteleosts. This phylogeny was then complemented with the detected genomic abundances in the considered species (Figure 4). We find that most highly abundant elements fall into the Minos-/Bari-like clade (Figure 4 clade A), such as the two most abundant elements of *U. pygmaea* (upygm-1-8, upygm-1-14), *E. masquinongy* (eluci-6-168), and *N. hubbsi* (eluci-4-461). Other more abundant lineages are passport (Figure 4 clade B and D), with the most abundant element eluci-1-0 in *E. lucius*, and frog prince (Figure 4 clade C, upygm-1-69, eluci-6-2394).

**Fig.4.**
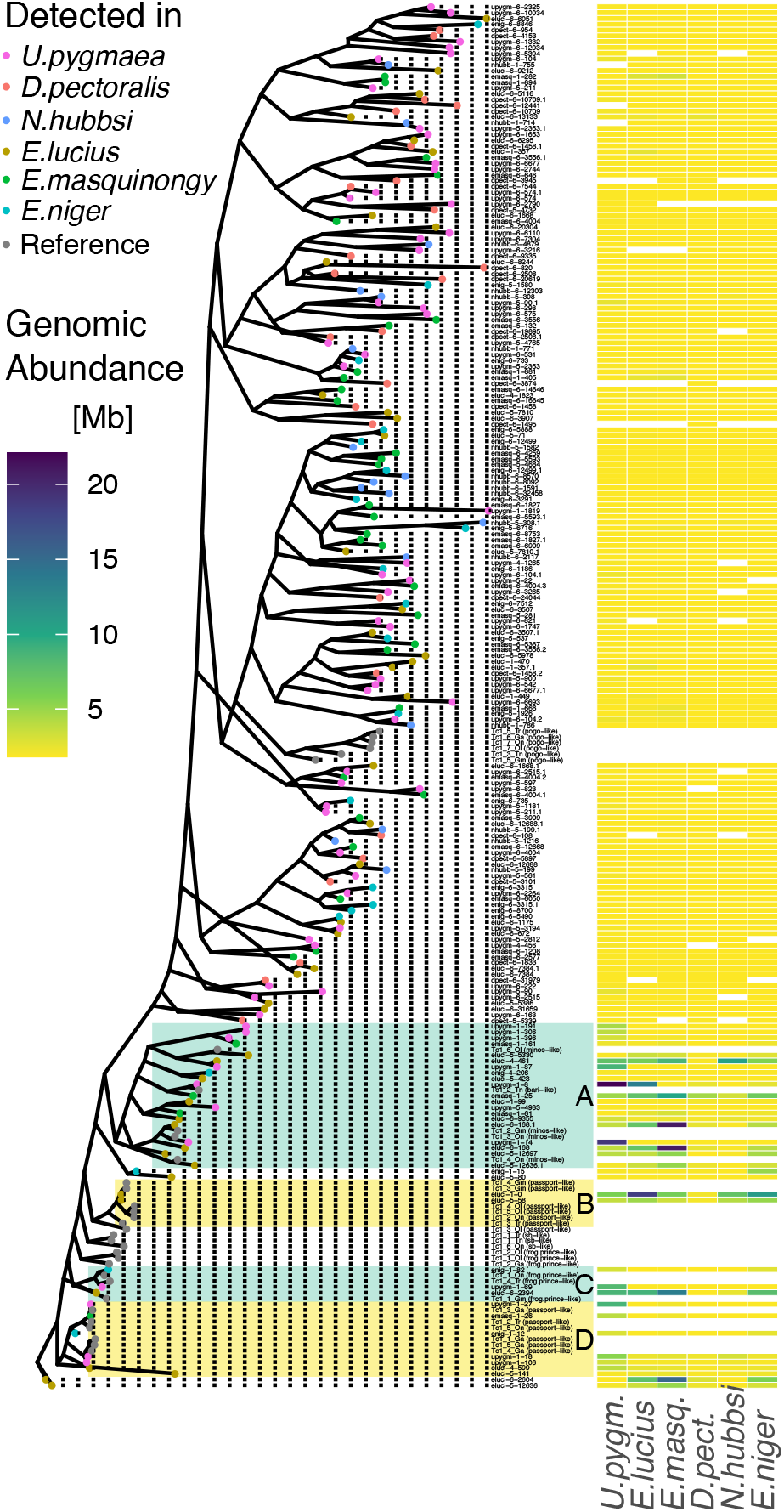
Phylogenetic relationship (left) and genomic abundance (right) of Tc1/mariner superfamily DNA transposons across six Esociformes species. The phylogenetic tree is based on transposase sequences from 206 repeats combined with 33 previously described(Gao et al. 2017) reference elements from six lineages (Pogo, Passport, SB, Frog Prince, Minos, and Bari). Clades containing most abundant elements are marked A-D. Genomic abundance values in Mb are based on full length repeat sequences, where elements not detected in the respective species are shown in white.

Finally, we performed a manual curation of the 30 most abundant repeat sequences found in *U. pygmaea*, contributing a combined 101.87 Mb to the genome assembly to test the extent to which the unclassified repeat sequence expansion can also be attributed to transposable element activity. We find TE-related structural features in 13 repeats, accounting for 48 Mb, in addition to a highly abundant tandem repeat and a satellite sequence (Table S6). With 11 out of 13 TE-related sequences, the vast majority shows DNA transposon features. In general, we observe a consistent pattern of increased abundance across species in a small group of Tc1/mariner elements mirroring the pattern of total genomic repeat abundance in the respective species.

### Fluorescence in Situ Hybridization

Studied pike species mostly exhibit karyotypes composed of 50 mono-armed chromosomes (FN = 50). In turn, *Umbra limi* possesses only 22 metacentric and submetacentric chromosomes (FN = 44), gradually decreasing in size. Fluorescence in Situ hybridization (FISH) with PNA telomere probe revealed hybridization signals only at the very ends of all chromosomes in *E. lucius* and *E. cisalpinus*. In *Umbra limi*, apart from the terminal location of the FISH telomeric signals observed on all chromosomes, up to seven chromosomes exhibited interstitial telomeric sites (ITSs) in the pericentromeric locations. In four of these chromosomes, ITSs overlapped with DAPI positive regions. DAPI positive signals occurred on the q-arms below the pericentromeric regions and did not correspond with the ITSs for up to three other chromosomes (Fig. 5A). The 5S rDNA probe hybridized to three sites in a pericentromeric location on two metacentric chromosomes and centromeric location on one submetacentric chromosome in *U. limi* (Fig. 5B). All 5S rDNA sites were DAPI-positive.

**Fig.5.**
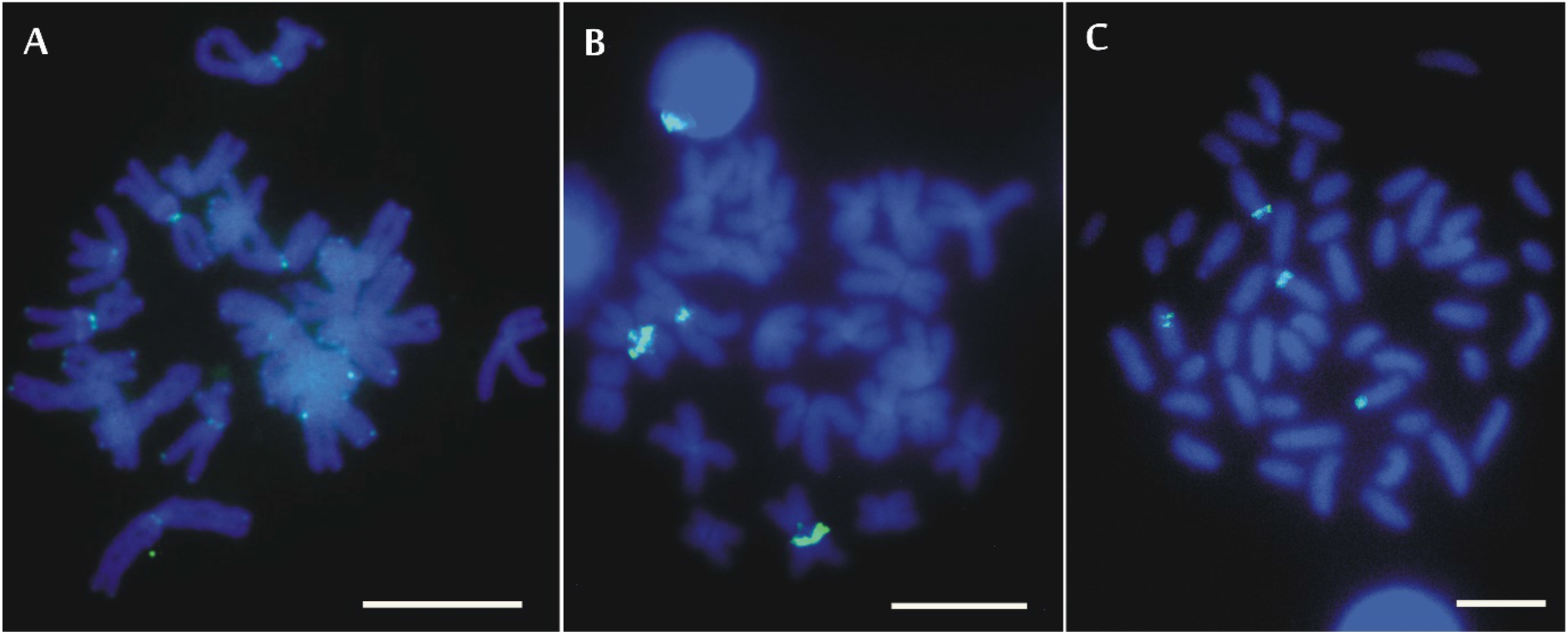
FISH with telomeric probes and 5S rDNA. Chromosomes of *Umbra limi* after FISH with telomeric probe (A) and 5S rDNA probe (B) and of *Esox niger* after FISH with 5S rDNA probe (C). All chromosomes are counterstained with DAPI. The FISH with a telomeric probe in *U. limi* yielded stronger interstitial telomeric signals than the terminal ones. Bar = 10μm.

### Molecular analysis of 5S rDNA genomic fraction

The slot blot quantification of 5S rRNA genes indicates approximately 2,000 5S rDNA copies per *Umbra* genome with a slightly higher copy number in *U. limi* compared to *U. krameri*. *Umbra* thus has about ten times lower copy number of 5S rRNA than both *Esox* species analyzed so far (Fig. 6 A, B). These copies are arranged in tandems in both *Umbra* as well as in *Esox* indicated by ladders of bands after the digestion with MspI restriction enzyme (not sensitive to CG methylation) (Fig. 6 C). Increased ladder length for both *Umbra* species reveals greater sequence divergence of MspI (CCGG) sites compared to *Esox*. Digestion of DNAs with methylation-sensitive HpaII isoschizomere resulted in hybridization signals in high-molecular-weight regions with little to no ladders (Fig. 6 C), which is consistent with heavy methylation of internal Cs within the CCGG motifs.

**Fig.6.**
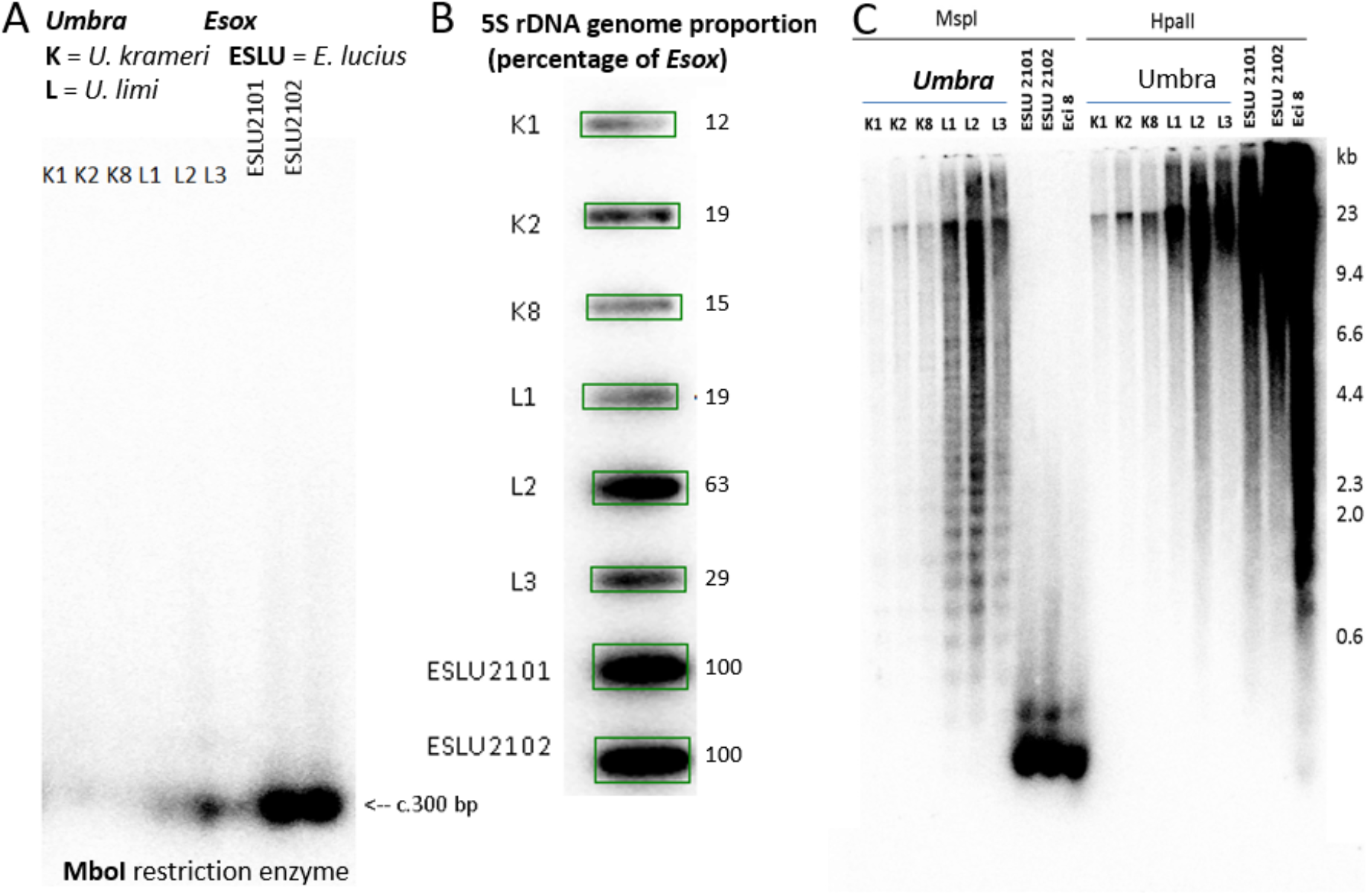
Genomic analysis of *Umbra* and *Esox* 5S rDNA. (A) Southern blot hybridization. (B) Slot blot hybridization. Probe – 5S rDNA insert from the *E. cisalpinus* clone a, K - *U. krameri*, L = *U. limi*, ESLU - *E. lucius*, (C) DNA methylation analysis of 5SrDNA in *Umbra* and *Esox. Hpa*II is a methylation-sensitive isoschizomere of *Msp*I. Probe – 5S rDNA insert from the *E. cisalpinus* (Eci) clone a. ESLU – *Esox lucius*, K – *Umbra krameri*, L – *Umbra limi*.

## Discussion

The sizes of eukaryotic genomes vary substantially with teleosts being no exception. The range of described cases of whole genome duplication (WGD) events amongst teleosts make this mechanism the main suspect when observing significant genome size increases. In contrast, we find that the genome expansion in the genus *Umbra* is not caused by a WGD but transposable element activity, and that extensive chromosome fusion events have occurred in some *Umbra* species. In the following, we discuss what sets these inconspicuous members of the order Esociformes apart from their sister lineage from an ecological and evolutionary point of view.

### Genome expansion and chromosome fusions in Umbra sp

Our results demonstrate that the European mudminnow species *U. krameri* underwent a genome expansion similar to the two North American *Umbra* sister species. This genome expansion is thus a common trait of the entire genus and can be localized in time prior to the split of the American and European *Umbra* lineages, setting the *Umbridae* family apart from their sister lineages *Esox*, *Dallia*, and *Novumbra* (Marić et al. 2017; Gregory 2020). The European species *U. krameri* retained the *Umbra* ancestral chromosome number (2n = FN = 44), which is even lower than the most common teleost ancestral 2n ≈ 50 (Mank & Avise 2006; Sacerdot et al. 2018) chromosome number and might indicate an overall tendency towards chromosome number reduction in the genus *Umbra*. Paradoxically, the genome expansion in *U. limi* and *U. pygmaea* is accompanied by a further reduction in chromosome counts (2n = 22) while maintaining the same number of chromosome arms (FN = 44). This reduction in the number of chromosomes without changes to the number of chromosome arms, as confirmed for *U. limi*, is consistent with Robertsonian fusion events. Accordingly, up to seven out of 22 bi-armed chromosomes of *U. limi* exhibit interstitial telomeric sites (ITSs), which presumably are remnants of these fusions. Similar occurrences of telomeric DNA at non-telomeric sites as relics of chromosomal rearrangements including centric fusions (Robertsonian translocations), tandem fusions, and inversions (Ocalewicz 2013) have been observed at pericentromeric locations of fused mammalian and non-mammalian chromosomes (Ocalewicz et al. 2013; Meyne et al. 1990). Breaks in telomer-adjacent chromatin regions before the fusion event may explain the absence of telomeric repeat sequences at the remaining fusion sites (Nanda et al. 1995; Garagna et al. 1995). Furthermore, ITSs may undergo gradual loss and degeneration, leading to progressive shortening. So while FISH is currently the best approach to explore ITSs *in situ*, some interstitially located telomeric DNA repeats might be too short for detection (Slijepcevic 1998).

The low chromosome count of the North American *Umbra* species (2n = 22) makes them part of a small group of fishes. Among about 32,000 fish species (Nelson et al. 2016), there are only thirteen fish species with 2n < 22 chromosomes described to date, made up mainly of Cyprinodontiformes (six in Nothobranchiidae, one in Rivulidae) ranging between 2n = 16 - 20. The lowest chromosome counts were described for the marine teleost species spark anglemouth (*Sigmops bathyphilus,* Stomiiformes, Gonostomatidae) (Post 1974) with 2n = 12 and the freshwater teleost species chocolate gourami (*Sphaerichthys osphromenoides*, Osphronemidae, Anabantiformes) (Calton & Denton 1974; Koref-Santibanez & Paepke 1994; Arai 2011) with 2n = 16. Interestingly, the *S. bathyphilus* belongs to the order Stomiiformes, a member of the superorder Osmeromorpha that split after the earlier divergence of the superorder Protacanthopterygii (i.e. Esociformes and Salmoniformes) (Nelson et al. 2016).

### Interplay between cytogenomic and ecological traits

The chromosome number of a species can be linked to its ecological traits. Specialized fishes were demonstrated to have smaller genomes than more generalized species within and across lineages (Hinegardner & Rosen 1972; Hardie & Hebert 2004). This constraint may be extended to the chromosome number and genome size in specialized species (Hardie & Hebert 2004), or constraints on genome size linked to the heightened developmental complexity of specialized species (Gregory 2002). Similar principles have been proposed for mammals (Qumsiyeh 1994) where species with higher 2n or FN have a higher recombination rate because proper disjunction requires at least one chiasma per arm (Jensen-Seaman et al. 2004). Increased recombination not only leads to increased genetic variability, thereby allowing the utilization of a wider niche (Qumsiyeh 1994), but also a genome size reduction (Nam & Ellegren 2012). On the other hand, decreased recombination favors the fixation of new mutations and thereby potential speciation (Qumsiyeh 1994). Furthermore, decreased recombination can cause the genome to expand due to the accumulation of transposon insertions (Dolgin & Charlesworth 2008). The interaction between transposon activity and recombination rate may create positive feedback, where the accumulation of transposon insertion sites contributes to further suppress recombination (Kent et al. 2017; Fedoroff 2012). The positive correlation between recombination rate and effective population size (Mugal et al. 2015) requires the consideration of the ecology of *Umbra* and *Esox*. Both genera’s species belong to freshwater fishes at northern latitudes and hence are affected by bottlenecks and reduced effective population sizes within refugia during glaciation periods (Bernatchez & Wilson 1998). Low levels of genetic diversity in pikes have been described extensively (Skog et al. 2014) and may explain the extreme 5S rDNA amplification through genetic drift (Symonová et al. 2017).

Genomic differences found among mudminnows can hardly be related to variable ecological traits within this group, as the ecology of mudminnows appears surprisingly similar. The American central mudminnow (*U. limii*) has been termed a “habitat specialist and resource generalist” (Martin-Bergmann & Gee 1985), referring to the densely vegetated and often overgrown and deoxygenated swampy habitats in which this species is found. Accessory air-breathing capabilities allow mudminnows to use such marginal habitats in which no, or only a few other fish species can be found. Otherwise, the central mudminnow uses a wide variety of small invertebrate prey. This brief ecological characterization can be extended to all other mudminnows (*sensu lato*), including *Novumbra* and *Dallia*. Despite their evolutionary differentiation, all mudminnows are found in such characteristic habitat types (Kuehne & Olden 2014) which might be related to their low competitive abilities (Wanzenböck & Spindler 1995; Sehr & Keckeis 2017). All mudminnows possess air-breathing accessory capabilities and use a wide variety of small invertebrate prey (Kuehne & Olden 2014). An additional peculiarity of this group of fishes is the parental care behavior, which become territorial during spawning time guarding the eggs. In summary, most ecological traits of mudminnows are convergent or stable despite their divergence in phylogeny and the differences in genomics found in our study.

In contrast, pikes are found in a wide variety of habitats, densely overgrown lentic waters to open waters in large lakes or even brackish waters and riverine habitats. Furthermore, they are rather specialized piscivorous during juvenile and adult life stages. Conversely, they might be called “habitat generalists and resource specialists.” Whether such ecological differences may be related to genomic differences between mudminnows and pikes remains open.

Next to potentially ecological influences, there are also purely genomic factors driving genome size evolution. A common cause of genome size expansion is a WGD event (polyploidy) or repeat expansion due to transposon or satellite DNA. In the *Umbra* genus, however, the genome expansion was accompanied by a reduction in chromosome number, which suggests two scenarios.

i. The *Umbra* lineage has experienced a rapid chromosome loss (disploidy) following an additional WGD. Evidence from plants suggests that repeated WGD cycles with subsequent chromosome loss may generate karyotypes with few chromosomes (Dodsworth et al. 2016). However, our comparative genome and transcriptome analyses do not support the existence of additional ohnologs in *Umbra pygmaea*. The observed number of duplicated protein-coding genes in the genome assemblies of *U. pygmaea* and its sister lineage *Esox lucius* was similar, excluding WGD as cause of genome expansion in *Umbra*. While the completeness of the employed transcriptomes and genomes is relatively low, it is not plausible that specifically ohnologs are disproportionately missing in both cases.
ii. Alternatively, the expansion of repeat sequences could explain the increased genome size. Evidence of such a repeat expansion was found in a sister lineage. The two species *E. lucius* and *E. cisalpinus* show an extreme amplification of 5S rDNA across their chromosomes (Symonová et al. 2017) while retaining their genome size and chromosome counts. This massive expansion of tandem repeats by several orders of magnitude must have happened over a short evolutionary time, as the related *E. niger* still features a low number of 5S rDNA copies ancestral to the Esocidae lineage (Kovařík, unpublished data). While the *Umbra* lineage shows teleost-typical(Sochorová et al. 2018) 5S rDNA copy numbers, our comparative genome analysis of the *U. pygmaea* genome confirms a significantly elevated interspersed repeat content of 64%, which is the result of recent transposon expansion peak driven by DNA transposon activity in combination with unclassified repeat sequences. This peak is not only consistent with the pervasive DNA transposon activity particularly in *E. lucius*, *E. masquinongy*, and *E. niger,* but also with the generally high abundance of DNA transposons in most fish genomes (Shao et al. 2019). Manual curation of the top 30 highly abundant unclassified sequences reveals DNA transposon features, further supporting the observed DNA transposon expansion. The Alaska blackfish *D. pectoralis* constitutes an exception as it appears in the process of repeat expansion driven by LINE elements, rRNA, tRNA, and unclassified elements. Importantly, genome size variation is the net result of sequence gain and loss (Petrov et al. 2000) allowing for the possibility of sequence gain by TE expansion in *U. pygmaea*, or alternatively sequence loss of various extent in the other species. As the expansion peaks observed in all species except *N. hubbsi* appear to be relatively recent at divergences below ten percent, a sequence gain scenario is more plausible. As all but the *E. lucius* genome assembly are highly fragmented and lack the completeness of reference genome assemblies, it is however likely that the presented results underestimate the total repeat content, particularly for long repeat sequences. This fragmentation is also a likely explanation for the high abundance of unclassified repeat sequences in *U. pygmaea*. Furthermore, it is not possible to locate chromosomal fusion sites or potential duplications in the *U. pygmaea* assembly. Further analyses based on a platinum standard long read based reference assemblies and an extensively curated library of repeat sequences are required to address these questions.

Here, we show that all members of the *Umbra* genus feature a significantly expanded genome, placing the expansion event prior to the separation between the European and North American species. However, only the North American species experienced Robertsonian chromosome fusion events, reducing their chromosome number significantly. Taken together, our findings show that a recent activity peak mainly of Tc1/mariner DNA transposons significantly increased the size of the *U. pygmaea* genome. We furthermore find evidence of an ongoing repeat-driven genome expansion in the Alaska blackfish *Dallia pectoralis*. This makes the family Umbridae and the entire order Esociformes an attractive model system to study exclusively repeat-driven genome expansion and its potential role in speciation.

## Material and Methods

### Fish sampling

Ten individuals of *Umbra krameri* were descendants from individuals sampled between 1993 - 1997 from “Fadenbach” between Orth and Eckartsau (Danube Wetlands, National Park in Lower Austria) within a nature conservation project of the provincial government of Lower Austria (Wanzenböck & Spindler 1995). Ten individuals of *Umbra limi* and ten individuals of *Esox niger* were sampled in Quebec, Canada, in autumn 2017. A local professional fisherman provided samples of two individuals of *E. lucius* at Lake Mondsee in 2016. Further specimens of *E. lucius* and *E. cisalpinus* were analyzed in our earlier study (Symonová et al. 2017). This included 28 eight-month-old specimens (12 males, nine females, and one unsexed) from the local fish farm (Olsztyn, Poland) sampled for cytogenetic analyses.

### Genome size determination by flow cytometry

Ethanol fixed red blood cells (RBC) of *Umbra krameri* and *Esox lucius*, respectively, were used for genome size determination following (Lamatsch et al. 2000) with chicken RBC as internal standard (C-value 1.25 pg per nucleus). We determined genome size by flow cytometry using the Attune™ NxT Acoustic Focusing Cytometer (Thermo Fisher Scientific, Vienna, Austria). We applied instrument settings optimized using the AttuneR Cytometric Software. We used forward scatter (FSC) and VL1 violet fluorescence (405 nm excitation, bandpass filter 440/50 nm) as triggers for all measurements. We analyzed 20.000 cells or sample volumes of 0.1 mL at a flow rate of 0.025 mL/min. DAPI is an AT-specific stain, which can yield inaccurate results when the AT/GC content of the sample and internal standard differ significantly. Chicken and *E. lucius* RBC, however, show a very similar AT-content of 57.73%, ^20,^ and 56.8%, (Borisova et al. 1974), respectively, suggesting that the underestimation of the total amount of DNA is negligible. Records on genome size come from the www.genomesize.com database (Gregory 2020) and the cited literature (Table 1).

### Comparative transcriptome analysis

We analyzed 24 *de novo* transcriptome assemblies of the PhyloFish database (Pasquier et al. 2016): bowfin (*Amia calva*), gar (*Lepisosteus oculatus*), European eel (*Anguilla anguilla*), butterflyfish (*Pantodon buchholzi*), arowana (*Osteoglossum bicirrhosum*), elephantnose fish (*Gnathonemus petersi*), Allis shad (*Alosa alosa*), zebrafish (*Danio rerio*), panga (*Pangasianodon hypophthalmus*), a black ghost (*Apteronotus albifrons*), cave Mexican tetra (*Astyanax mexicanus*), surface Mexican tetra (*Astyanax mexicanus*), Northern pike (*Esox lucius*), Eastern mudminnow (*Umbra pygmaea*), grayling (*Thymallus thymallus*), European whitefish (*Coregonus lavaretus*), American (Lake) whitefish (*Coregonus clupeaformis*), brown trout (*Salmo trutta*), rainbow trout (*Oncorhynchus mykiss*), brook trout (*Salvelinus fontinalis*), Ayu (*Plecoglossus altivelis*), Atlantic cod (*Gadus morhua*), medaka (*Oryzias latipes*), and European perch (*Perca fluviatilis*). The completeness and duplication were assessed with BUSCO v4.1.4 (Simão et al. 2015) using the dataset actinopterygii_odb10 containing 3640 genes (Table S1). For a subset of 10 species, reference genome assemblies and structural genome annotations are available at the NCBI reference sequence database (Table S2). For this subset, *de novo* transcriptome assembly completeness and duplication were compared to reference-genome-derived transcriptomes using identical BUSCO settings. Gene counts were separately compared for each of the five gene categories (complete, complete-single-copy, complete-duplicated, fragmented, missing) using the coefficient of determination (Fig. S1). A phylogenetic profile matrix was assembled representing the number of orthologous copies detected for each of the 3640 BUSCO genes across 24 transcriptome assemblies. Hierarchical clustering of the phylogenetic profile matrix was performed using Manhattan distance and the “ward.D2” method implemented in R v3.6.1 to obtain five main gene clusters and 26 sub-clusters of main cluster III.

### Comparative genome analysis

The five published(Pan et al. 2021) short-read based scaffold-level genome assemblies of the Esocidae species *Umbra pygmaea*, *Dallia pectoralis*, *Novumbra hubbsi*, *Esox niger*, and *Esox masquinongy,* as well as a chromosome-scale reference genome assembly of *Esox lucius,* were used for a comparative genome analysis (see Table S4 for accession numbers). For each of the six genomes, a *de novo* repeat library was generated using RepeatModeler 2.0.1, which yielded between 2,547 and 4,983 sequences per species. These libraries were then combined and clustered using usearch v11.0.667 with 80% similarity cutoff to obtain a single non-redundant repeat library for the order Esociformes, which yielded a total of 14,856 sequences. Censor 4.2.29 was used to identify *de novo* sequences in addition to the RepeatModeler classification and annotation with RepeatMasker and RepBase 26.01. This library was then used as a reference to annotate repeat sequences with RepeatMasker 4.1.1. Repeats containing known TE coding domains were identified by predicting open reading frames using Emboss v6.6.0.0 and comparing the resulting sequences against the transposase protein sequence collection provided with LTR_retriever(Ou & Jiang 2018) via blast. Requiring hits longer than 150 amino acids and with e-value < 1e^−10^ yielded 116 hits in LINE elements and 206 hits in sequences not classified as LINE elements, the latter of which were retained for further analysis. This procedure was also used to extract 33 transposase sequences of six previously described neoteleost Tc1/mariner lineages(Gao et al. 2017). All 239 sequences were then aligned with Clustal Omega v1.2.4, and a maximum likelihood phylogenetic tree was calculated with RAxML 8.2.12 using a variable time amino acid substitution model with gamma distributed rate heterogeneity. Analysis and visualization of all data was performed using R 3.6.1. The thirty most abundant *Umbra pygmaea* repeat sequences of the *de novo* library which did not show significant similarity with known TEs were selected for further manual curation. Firstly, we tested if the unclassified sequences include local context sequence in addition to the core repetitive tract. To that end, the repetitive sequence was used as a query in a Blastn (Altschul et al. 1997) search against the *Umbra pygmaea* genome assembly. Two or more matching loci located on different contigs/scaffolds were extracted including 10 Kbp up- and downstream of the matching sequence. These hit sequences were then compared to each other in a dot plot analysis using the dotter (Sonnhammer & Durbin 1995), allowing the exact definition of the entire repetitive element. Complete elements were then analyzed for similarity with known TEs as defined in the latest version of RepBase (Bao et al. 2015) and/or for the presence of structural features typical of known TE elements such as the inverted repeats of DNA TEs. This allowed the association of 13 elements to a TE class including 2 cases with only diagnostic structural features (Table S6). In addition, one satellite and one tandem repeat sequence were identified, leaving a total of 14 sequences unclassified. The curated sequences are provided at https://github.com/roblehmann/umbridae_genome_expansion.

### Chromosome analyses and fluorescence in situ hybridization

Telomeric DNA repeats on chromosomes of *Umbra* and *Esox* species were detected by FISH, using a telomere PNA (peptide nucleic acid) FISH Kit/FITC (DAKO, Denmark) according to the manufacturer’s protocol. Slides with the metaphase spreads were washed with buffer (Tris-buffered saline, pH 7.5) for 2 min, immersed in 3.7 % formaldehyde in 1× TBS for 2 min, washed twice in TBS for 5 min each, and treated with the Pre-Treatment solution including Proteinase K (DAKO) for 10 min. Afterward, slides were washed twice in TBS buffer for 5 min each, dehydrated through a cold (−20 °C) ethanol series (70 %, 85 %, 96 %) for 1 min, and air-dried at room temperature. Ten μl of FITC PNA telomere probe mix (DAKO) was dropped on the prepared slides and covered with the coverslip. Chromosomal DNA was denatured at 85° C for 5 minutes under the coverslip in the presence of the PNA probe. The hybridization reaction took place in the darkness at room temperature for 90 minutes. After hybridization, the coverslips were gently removed by immersion of slides in the Rinse Solution (DAKO) for 1 min. Slides were then washed in the Wash Solution (DAKO) for 5 min at 65°C and dehydrated by immersion through a series of cold ethanol washes of 70%, 85%, 96% for 1 min each and air-dried at room temperature. For the counterstaining, chromosomes were mounted in the VECTASHIELD Antifade Mounting Medium containing DAPI (Vector Laboratories, USA).

Metaphase plates were analyzed under a Zeiss Axio Imager A1 microscope equipped with a fluorescent lamp and a digital camera. The images were electronically processed using the Band View/FISH View software (Applied Spectral Imaging, Galiliee, Israel). FISH with 5S rDNA probe was performed according to (Fujiwara et al. 1998) with slight modification (Kirtiklis et al. 2014). A 5S rDNA probe was obtained via PCR with forward primer 5S-1: 5’-TACGCC CGATCT CGT CCG ATC - 3’ and reverse primer 5S-2: 5’-CAG GCTGGT ATG GCC GTA AGC - 3’ (Pendas et al. 1994). The PCR reaction was carried out in a 50 μl reaction volume containing 1.25 U GoTaq Flexi DNA Polymerase (Promega, USA), 10 μl of 5X Flexi Reaction Buffer (Promega, USA), 100 μM of each dNTP (Promega, USA), 3 mM of MgCl_2_ (Promega, USA), 10 pM of each primer, 2 μl of DNA template, and nuclease-free water. PCR product, obtained after 30 cycles of amplification and annealing at 55 °C, was purified using the GeneElute PCR Clean-Up Kit (Sigma-Aldrich, USA), then labeled with biotin-16-dUTP (Roche, Germany) by nick-translation method (Roche, Switzerland). *In situ* hybridization with 150 ng of rDNA probe per slide was performed with RNase-pre-treated and formamide-denaturated chromosome slides. A post-hybridization wash was performed at 37°C for 20 min. Chromosome slides were subjected to the detection with avidin-FITC (Roche, Basel, Switzerland) and then counterstained with DAPI in VECTASHIELD Antifade Mounting Medium (Vector Laboratories, USA). At least 15 metaphase chromosome spreads from each specimen were analyzed using a Nikon Eclipse 90i (Nikon, Japan) microscope equipped with epi-fluorescence. Pictures were acquired using a monochromatic ProgRes MFcool camera (Jenoptic, Germany) controlled by a Lucia software ver. 2.0 (Laboratory Imaging, Czech Republic). Post-processing elaboration of all the pictures was made based on CorelDRAW Graphics Suite 11 (Corel Corporation, Canada).

### Southern and slot blot hybridizations of 5S rDNA

The procedure followed the protocol described by (Koukalova et al. 2010). The 5S rDNA probe was a 243 bp insert of the 5S_Eci_a clone (GenBank KX965716) from *E. cisalpinus*. The plasmid insert was amplified and labeled with the 32P-dCTP (DekaPrime kit, Fermentas, Lithuania). The probe was hybridized at high stringency conditions (washing 2x SSC, 0.1% SDS followed by 0.1xSSC, 0.1% SDS at 65 °C). The hybridization signals were visualized by Phosphor imaging (Typhoon 9410, GE Healthcare, PA, USA), and signals were quantified using ImageQuant software (GE Healthcare, PA, USA). The copy number of 5S rDNA genes was estimated using slot blot hybridization. Briefly, the DNA concentration was estimated spectrophotometrically at OD260nm using a Nanodrop 3300 fluorospectrometer (Thermo Fisher Scientific, USA). Concentrations were verified by the electrophoresis in agarose gels using dilutions of lambda DNA as standards. The three dilutions of genomic DNA (100, 50, and 25 ng), together with serial dilutions of unlabelled plasmid inserts corresponding to the 5S monomers (GenBank KX965715-6). Each aliquot was denatured in 0.4 M NaOH and blotted onto a positively charged Nylon membrane (Hybond XC, GE Healthcare, USA) using a vacuum slot blotter (Schleicher-Schuell, Germany). The probe and the hybridization conditions and visualization of signals were the same as described above.

### DNA methylation analysis of 5S rDNA

Purified genomic DNA samples of *U. krameri* (K1 - K3), *U. limi* (L1 - L3), and from *E. lucius* (2 individuals as a control) were digested with methylation-sensitive *Hpa*II (sensitive to CG methylation) and its methylation-insensitive *Msp*I isoschizomere (both enzymes are cutting at CCGG). The restriction fragments were hybridized on blots with the alpha[P^32^]dCTP-labelled 5S rDNA probe. Control of digestion efficiency was carried out by spiking the *Esox* genomic DNA with a non-methylated plasmid DNA (pBluescript, Stratagen) and subsequent hybridization with a plasmid probe. Both *Msp*I and *Hpa*II enzymes yielded expected restriction fragments (not shown).

## Author Contributions

RS conceptualized and supervised the study and provided the cytogenomic context; RS, AK, RL, and DKL methodized the study and wrote the original draft; RL analyzed fish transcriptomes and genomes; RL and AZ analyzed transposable elements; AK performed southern blot and slot blot hybridizations; DKL performed the flow cytometry genome size determination; JW provided the ecological and phylogenetic context of the study and specimens for analyses; LB organized sampling, provided specimens for analyses and reviewed the manuscript; KO, LK performed the cytogenetic analyses; RL, RS, DKL, KO, LK, JW, JT wrote, reviewed and edited the study; RS acquired funding for the study.

## Acknowledgments

This study was supported by the Tyrolean funds project with contract Nr. UNI-0404/2015 to RS and co-funded by the Erasmus+ program of the European Union with contract Nr. 2019-1-CZ01-KA203-061433 to RS and DKL. The authors are also grateful to the ‘Excelence projekt PřF UHK 2209/2018’ and the Czech Science Foundation (19-03442S) for financial support.

We acknowledge Petr Ráb for discussion on genome size in the *Umbra* genus, Guillaume Côté for fishing *Umbra* and *Esox* in Québec, and we also acknowledge Maria Pichler of UIBK for her technical support.

## Data Availability

The *de novo* repeat library and manually curated sequences are available via https://github.com/roblehmann/umbridae_genome_expansion

