## Supplementary Figures for "DNA transposon expansion is associated with genome size increase in mudminnows"

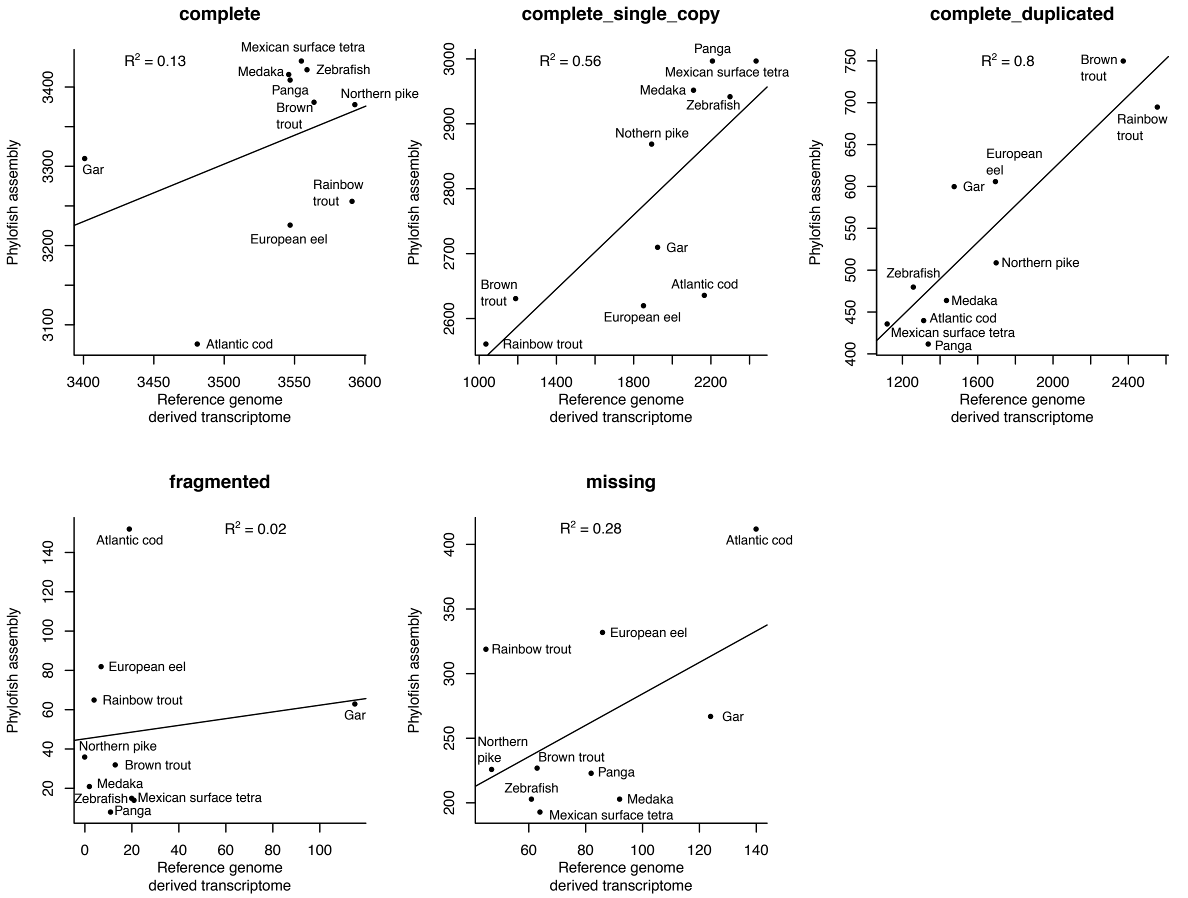


**Fig.S1**. Comparison of BUSCO gene counts between *de novo* transcriptome assemblies and transcriptomes derived from reference genome assemblies of the same species. The coefficient of determination between *de-novo* transcriptome assembly values and reference genome transcriptome values is show in each panel.

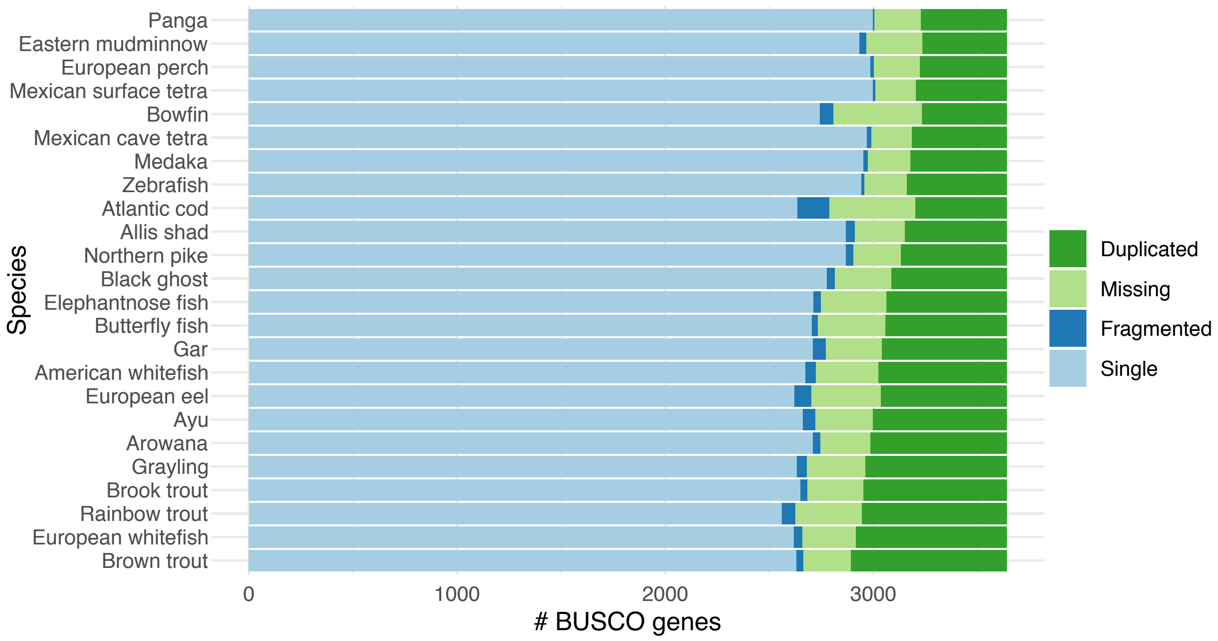


**Fig.S2.** Comparison of Actinopterygii-specific universal single-copy orthologous gene counts of *de-novo* transcriptome assemblies for 24 fish species. Species are ordered by ratio of complete to duplicated gene count.

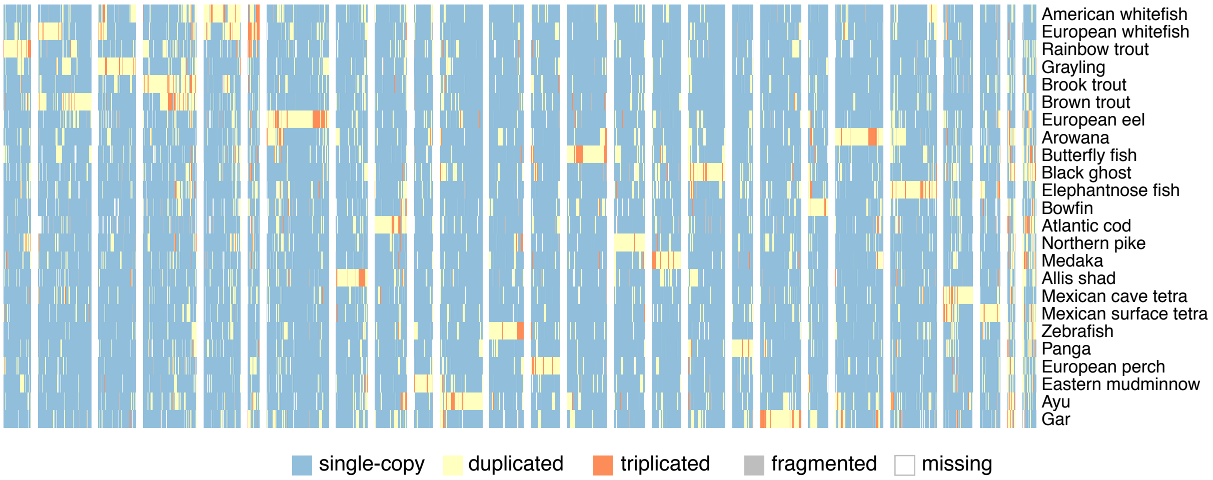


**Fig.S3**. Sub-clustering of phylogenetic profiles of genes in cluster III. For the number of genes in each cluster, refer to Table S3.

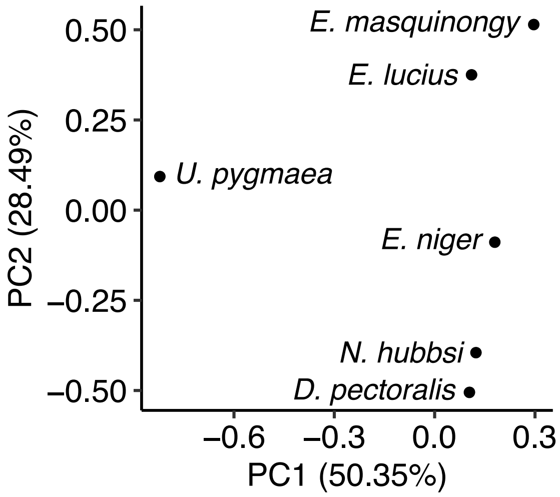


**Fig.S4.** Principal component analysis of repetitive element content for six Esociformes species.
